# Improved workflow for untargeted metabolomics and NMR analysis of intracellular and extracellular metabolites isolated from Gram positive and Gram negative bacteria

**DOI:** 10.1101/2023.11.24.568533

**Authors:** Dean Frawley, Trinidad Velasco-Torrijos, Fiona Walsh

## Abstract

Metabolomics and Nuclear Magnetic Resonance (NMR) spectroscopy have proven to be useful for characterising key metabolome fluctuations in bacteria during stress responses to various environmental agents and antibiotics. However, a number of impediments to current workflows have led to the reduced use of these techniques in microbial research. In this study, we address these limitations and in response have developed a novel ^1^H NMR-based untargeted metabolomics workflow. This method is suitable for use with various bacterial species, reducing the workload in comparison to previously established workflows. Our protocol is simple and reproducible and allows for the isolation of both intracellular (IC) and extracellular (EC) metabolites simultaneously from both Gram (+) and Gram (-) species. This method has been shown to produce consistent results for the ESKAPE pathogens *Escherichia coli, Klebsiella pneumoniae, Enterococcus faecium* and *Staphylococcus aureus*. By using these data as a baseline, future studies involving a myriad of stress conditions can be compared to identify key metabolome differences in each species and to determine the mechanisms utilised by bacteria to respond to stress.

## 1. Introduction

Metabolomics is a broad term that describes techniques used to identify and quantify organic molecules like amino acids, carbohydrates, lipids, nucleic acids and primary and secondary metabolites that are generated by cells. Metabolomics has proven to be particularly useful as evidence suggests that various metabolic processes in bacteria may influence antibiotic susceptibility (Bush et al., 2011; Lobritz et al., 2015). Metabolomics is considered the endpoint of cellular stress-responses and is the omics method that is most closely related to the biological phenotype (Pan et al., 2021). Any changes observed at the metabolome level are often a result of perturbations in upstream genomic, transcriptomic or proteomic cascades. Thus, metabolomic studies are regularly performed with the aim of bridging the gap between the phenotype observed and various omics data, such as genomics (Tounta et al., 2021). Untargeted metabolomics aims to investigate all compounds isolated from a sample. It has been used to analyse global metabolome changes and to detect and quantify metabolites produced by bacteria in response to various stress conditions. These include oxidative stress, temperature changes, osmotic stress, pH perturbations and antibiotic stress, to name a few (Fiehn, 2002; Gaucher et al., 2020; Kurake et al., 2019; Patejko et al., 2017; Patti et al., 2012). By analysing the metabolic profiles of bacteria cultured in various growth conditions, this can provide information regarding the mechanisms utilised to respond to particular stress agents. In relation to antibiotic stress, this can aid in the identification of potential drug targets and can help elucidate the modes of action of various antibiotics.

Intracellular (IC) metabolomics is the study of metabolites that are contained within mechanical barriers, such as the cell membrane. The full array of IC compounds in a cell are collectively known as the endo-metabolome or metabolic fingerprint (Villas-Bôas et al., 2005). Extracellular (EC) metabolomics defines the study of metabolites that have been secreted by cells into their surrounding environment. The exo-metabolome, or metabolic footprint is the entire repertoire of metabolites found outside the microbial cell in the supernatant (Mashego et al., 2007). By combining exo-metabolome and endo-metabolome analysis, this can help provide a detailed view of microbial metabolism in response to a specific environmental condition or stress (Pinu and Villas-Boas, 2017). However, certain challenges exist when isolating both IC and EC metabolites. The process of isolating these compounds is extremely sensitive and many factors can influence the levels of metabolites, including light, temperature, pH, concentration of nutrients and metabolite interactions (Mapelli et al., 2008). Thus, techniques have to be performed to avoid dramatic changes in the metabolite profile. Once the desired growth conditions have been met for an experiment, the cells must be quickly removed from the growth medium and a quenching solution used to stop cell metabolism in order to reduce the risk of metabolite degradation (Patejko et al., 2017; Villas-Bôas and Bruheim, 2007). Bacteria are highly prone to leaking intracellular metabolites during the quenching process and thus it is essential to use quenching solutions and methods that limit damage to the cell membrane, resulting in reduced leakage and increased recovery and detection of metabolites (Pinu et al., 2017). The bacterial membrane is an effective barrier for the cell, helping to retain metabolites. However, this barrier can be disrupted by the use of organic solvents and detergents as these help dissolve lipids in the membrane. Thus, solvents and detergents such as methanol, ethanol, chloroform and sodium-lauryl sarcosinate are often used in metabolite extraction procedures to disrupt the cell membrane and to inhibit enzyme function to help increase metabolite recovery (Clark and Beard, 1979; Filip et al., 1973; Mielko et al., 2021).

Nuclear magnetic resonance (NMR) is a spectroscopic technique that has been utilised in a myriad of scientific disciplines. NMR can be used to unambiguously identify organic molecules and to provide information regarding their structure by detecting the energy transitions of atomic nuclei when in the presence of a magnetic field (Moco, 2022). Aside from studies on individual molecules, NMR can also be applied for the analysis of proteins, nucleic acids and metabolites isolated from cells, as well as metabolite-protein interactions (Moco, 2022; Vignoli et al., 2019). NMR offers significant advantages when compared to other technologies used for metabolomics research, such as mass spectrometry (MS)-based methods (Zhong et al., 2022). While MS-based methods enable the analysis of the molecular mass and formula of a compound, they only reveal some structural features. However, NMR, is capable of revealing the entire structure of a compound (Bruice, 2016). NMR measurements are also highly robust, stable and reproducible over months to years, assuming samples are stored correctly (Pinto et al., 2014; Ward et al., 2010). The reproducibility of NMR has allowed for the standardisation of metabolomics procedures in research and clinical settings (Emwas et al., 2015; Ward et al., 2010). Another advantage of NMR is that chromatographic methods are not necessary, meaning that samples are not in contact with the instrument, thus reducing the need for instrument cleaning and minimising contamination. Also, NMR is quantitative, which enables the detection of both relative and absolute concentrations for metabolites and it is capable of quantifying many metabolites simultaneously (Moco, 2022).

While NMR-based metabolomics is useful for researching bacterial metabolism, there are a number of limitations to current workflows. Protocols that are suitable for isolating both IC and EC metabolites simultaneously do exist and have been proven to be successful for singular species (Hoerr et al., 2016; Qiao et al., 2019). However, these protocols are scarce and in order to isolate both groups of metabolites, the extraction procedures are usually performed separately, leading to batch variation. It is also uncommon that metabolomics studies are performed concurrently on multiple groups of bacterial species. Consequently, metabolomics workflows are often restricted to individual species or individual types of metabolites. Lastly, the majority of studies are focused solely on the analysis of IC compounds and EC analysis is often excluded, commonly due to sugar contamination in ^1^H NMR spectra which is caused by carbohydrates present in microbial growth media (Yuan et al., 2018).

We aimed to establish an improved, simplified and reproducible workflow for untargeted metabolomics, applicable to both Gram (+) and Gram (-) bacteria. This workflow enables the isolation of both IC and EC metabolites from Gram (+) and Gram (-) bacteria simultaneously, followed by the processing and analysis of these metabolite samples with NMR and Chenomx software. This protocol allows for simplified metabolome characterisation in a range of bacterial species, reducing the workload in comparison with current NMR-based metabolomics workflows.

## 2. Methods

### 2.1. Strains and culture medium

The following strains were used for all experiments: *Escherichia coli* MG1655*, Klebsiella pneumoniae* NCTC418*, Enterococcus faecium* NCTC13169 and *Staphylococcus aureus* NCTC8325. For all metabolomics experiments, strains were cultured in a modified M9 minimal medium containing 1X M9 salts solution (M6030, Sigma Aldrich), 0.4% glucose, 2mM MgSO_4_, 0.1mM CaCl_2_ and 0.5% yeast extract.

### 2.2. Isolation of intracellular and extracellular metabolites

Isolated colonies for each strain were inoculated in triplicates of 5mL M9 minimal medium and were cultured overnight in a shaking incubator at 225 RPM and 37°C overnight. Each 5mL culture was diluted in a total of 100mL M9 minimal medium in a sterile 250mL flask. The absorbance at OD_600_ of each culture was recorded and the samples were incubated at 225 RPM and 37°C until the absorbance reached the mid-exponential phase (absorbance=0.35). Then, 10μL of sterile water was added to the samples to act as a control and the samples were incubated at 225 RPM and 37°C for 1 hour. After this incubation, each 100mL culture was centrifuged at 3,000 RPM for 10 minutes at 4°C.

To isolate extracellular compounds, 20mL of supernatant per sample was filtered through a 0.2μm filter into a 50mL Falcon tube and this was frozen with liquid nitrogen. These extracellular metabolite extracts were then lyophilised and stored at -80°C and the remaining 80mL supernatant was discarded. To isolate intracellular compounds, cells were quenched using previous protocols, (Aries and Cloninger, 2020), with some variations. The cells were quenched by resuspending the pellet in Methanol (800mL stored at -80°C) and ice-cold double distilled water (170mL). The resuspended sample was vortexed and added to a 2mL tube. The sample was sonicated in a water bath for 10 minutes. To isolate intracellular metabolites, 800mL chloroform (stored at -80°C) and 400mL of ice-cold water was then added and the sample was transferred to a 15mL Falcon tube. All samples were vortexed and left on ice for 15 minutes. Samples were then centrifuged for 20 minutes at 4°C at 3,000 RPM. The top aqueous methanol layer containing the intracellular metabolites was then pipetted into a new 2mL tube and this was dried in a SpeedVac. The dried extracts were stored at -80°C until proceeding with acetone precipitation.

### 2.3. Acetone precipitation

After drying all intracellular and extracellular samples, each sample was resuspended in 250μL ice-cold NMR-grade deuterium oxide (166301000, Thermo Scientific) and 1.25mL acetone (stored at -80°C) (Aries and Cloninger, 2020) and incubated at -80°C overnight. The next day, samples were thawed on ice and centrifuged at 2,000 RPM for 30 minutes at 4°C. The liquid was then transferred to new 2mL tubes and samples were dried in a SpeedVac. Dried samples were stored at -80°C.

### 2.4. Sodium periodate (NaIO_4_) treatment

NaIO_4_ treatments were performed, as according to (Yuan et al., 2018). Prior to NaIO_4_ treatment, all samples were equilibrated to room temperature. All intracellular metabolite samples were resuspended in 750μL deuterium oxide containing 1mM 3-(Trimethylsilyl)-1-propanesulfonic acid sodium salt (DSS) (178837, Sigma Aldrich) and 50mM NaIO_4_ (769517, Sigma Aldrich). All extracellular samples were resuspended in 750μL deuterium oxide containing 1mM DSS and 300mM NaIO_4_. Samples were left to incubate in the dark at room temperature for 4 hours. Glass Pasteur pipettes were then used to add samples into NMR tubes (ST550-7, Norell).

### 2.5. Data acquisition

1D ^1^H NMR spectra were recorded using a Bruker Ascend 500 spectrometer (500 Mhz) at 293K. ^1^H NMR spectra were acquired using the Bruker-automated pulse program ZG30 over 16 scans, in NMR-grade deuterium oxide (166301000, Thermo Scientific) with 3-(trimethylsilyl)-1-propanesulfonic acid sodium salt (DSS, 178837, Sigma Aldrich) as internal standard. ^1^H NMR spectra were Fourier transformed and processed using TopSpin Sofware (3.0). Chemical shifts were reported in ppm

### 2.6. NMR data analysis

All ^1^H NMR spectra were processed and profiled using the Chenomx NMR Suite software version 9.0 (Chenomx, Inc. Alberta, Canada). The Chenomx Processor module was used to manually correct the spectral data, while the Profiler module was used to identify and quantify metabolites and to generate a fitted spectrum for each sample. The concentrations of the metabolites were quantified by using 1mM DSS as an internal standard.

## 3. Results

### 3.1. Defining a growth medium and culture conditions for metabolomics experiments

In general, the process flow for metabolomics often involves sample culturing, separation of IC and EC metabolites, quenching of metabolism and isolation of IC compounds, followed by sample processing and analysis by suitable means (**Figure 1**). In order to optimise a suitable metabolomics protocol applicable to both Gram (+) and Gram (-) bacteria, we first had to define a minimal medium that could support the growth of both Gram (+) and Gram (-) bacteria. The test species used in the optimisation of the methods were *Escherichia coli* and *Klebsiella pneumoniae*, which are Gram (-) bacteria, as well as *Enterococcus faecium* and *Staphylococcus aureus*, which are Gram (+) species. We initially tested a base M9 formula (1X M9 salts, 0.4% glucose, 2mM MgSO_4_, 0.1mM CaCl_2_) for the minimal medium which is commonly used for bacterial cultivation. However, while we found this medium was sufficient for *E. coli* K-12 derivatives and *K. pneumoniae*, it was not suitable for the growth of *E. coli* DH5a strains or any of the Gram (+) species. We then decided to test a myriad of different supplements in the medium. It was found that no supplements tested other than yeast extract could sustain growth at a high enough level for each of the 4 species. Thus, yeast extract was chosen as the only supplement at a concentration of 0.5% to enable sufficient growth of both Gram (+) and Gram (-) species for this study.

**Figure 1.**
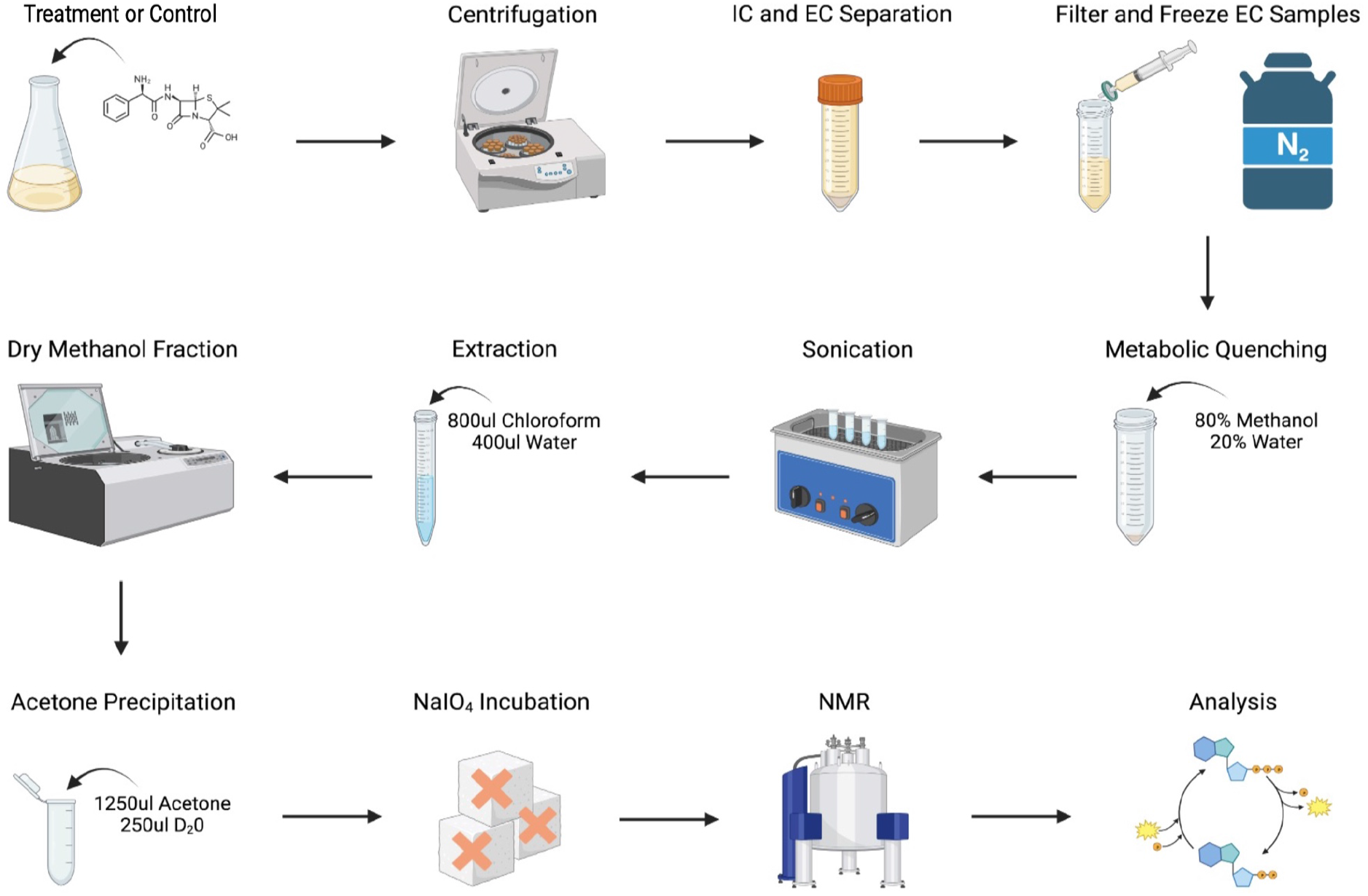
Workflow illustrating the main steps involved in isolating intracellular and extracellular metabolites from both Gram (+) and Gram (-) bacteria. After reaching an absorbance of 0.35, samples were subjected to a specific growth condition such as a control or treatment for 1 hour. For this work, all samples were subjected to a control condition which is 10μL sterile water. Samples were then centrifuged and the resulting cell pellet contains the intracellular (IC) metabolites, while the supernatant contains the extracellular (EC) metabolites. Then, 20mL of the supernatant was filtered and frozen using liquid nitrogen and was later lyophilised. The pellet undergoes metabolic quenching to inhibit metabolism and was sonicated to disrupt the cells. A chloroform:water solution was used to lyse the cells and release the IC metabolites. The isolated metabolites migrate to the aqueous methanol:water fraction which was dried using a SpeedVac. This extract was then acetone precipitated to solubilise and isolate the metabolites. NaIO_4_ incubations were then performed to remove sugars from the samples that can interfere with NMR analysis. ^1^H NMR 16 scan runs were then performed and the resulting raw spectra were processed using Chenomx v9.0 software, using the Processor and Profiler modules. Figure created using Biorender.

We then defined suitable growth conditions for the metabolomics experiments. After initially culturing strains overnight in M9 medium, the strains were sub-cultured and grown to an absorbance of 0.35. This signifies that the cells are in the mid-exponential phase and it is in this phase that the cells are most metabolically active (Qiao et al., 2019). Once this growth phase was reached, sterile water was added to the culture and samples were left to incubate for 1 hour to act as a control. This 1 hour timepoint was chosen as it has been shown previously that this gives the cells sufficient time to elicit a metabolic response (Ye et al., 2021). In future studies, the 1 hour water control will be compared alongside a 1 hour treatment. In this hour, the treatment should evoke a metabolic response from the cells without killing them, whereas the response to the water control should be minimal.

### 3.2. Optimisation of metabolic quenching and extraction of metabolites

After the 1 hour incubation, to separate the cell biomass from the culture medium, samples were centrifuged at low speeds and at 4°C to minimise cell damage. Then, 20mL of the supernatant containing the EC metabolites was filtered to remove any traces of cells (**Figure 1**). In order to inhibit any enzymatic reactions which may rapidly degrade metabolites, a process known as quenching is performed. (Patejko et al., 2017). There are numerous methods of quenching documented for various species, each with advantages and disadvantages (Pinu et al., 2017). For this protocol, the filtered supernatant is quickly quenched by freezing with liquid nitrogen. The frozen supernatants are then lyophilised to remove the medium and to concentrate the isolated EC metabolites. To quench the cells containing the IC metabolites, a cold methanol:water solution was used (**Figure 1**). Cold methanol solutions of various concentrations have been well documented in the literature and while there have been conflicting opinions regarding their use as quenching solvents, they are generally considered to be suitable for metabolomics experiments and have been shown to induce minimal cell leakage (Aries and Cloninger, 2020; de Koning and van Dam, 1992; Lei et al., 2020). These methods of quenching IC and EC metabolites in our protocol have been shown to be efficient for each of the 4 species tested.

In order to lyse the cells and release IC metabolites, a wide range of methods have been published for a myriad of species. It is often stated that various methods are specific for isolating certain types of compounds or are specific to a singular species, based on cell composition (Pinu et al., 2017). For this protocol, we decided to use a non-polar extraction solution consisting of cold chloroform and water (**Figure 1**). When added, this solvent lyses the cells and promotes isolation of IC metabolites. Non-polar solvents are often used to extract IC compounds from microbes. These solvents are capable of forming pores in the cell envelope, allowing for the release of IC metabolites which then migrate into the polar methanol solution (Aries and Cloninger, 2020; Pinu et al., 2017). The methanol solution can then be separated from the chloroform layer and dried in a SpeedVac to remove the liquid and concentrate the isolated IC metabolites. We have shown that this singular extraction solution is suitable for use with each of the 4 species tested. Once the IC and EC extracts are dried down, the metabolites are further purified by performing an acetone precipitation (**Figure 1**). Proteins and other macromolecules will precipitate as they are not soluble in acetone, while metabolites are soluble. Thus, after incubation in acetone and centrifugation, the solubilised metabolites can be isolated and dried down to remove the liquid and concentrate the metabolite extract.

### 3.3. Use of the NaIO_4_ oxidant for the removal of sugar contamination

Before running samples on an NMR spectrometer, sugars need to be removed as these can interfere with the NMR spectra. The 3.2-4.5 ppm region in a ^1^H NMR spectrum is often dominated by carbohydrates like glucose, sucrose and fructose, to name a few. These sugars exhibit high chemical similarity, leading to significant overlap of their NMR resonances (Yuan et al., 2018). This spectral crowding makes it difficult to detect individual carbohydrates, as well as non-carbohydrate metabolite signals in these regions of the spectrum, leading to a loss of valuable information. Unfortunately, it is not always possible to eliminate the use of sugars in metabolomics experiments as these carbohydrates are often needed in the growth media for many microbes. However, it has been shown that the oxidant sodium periodate (NaIO_4_) is mild and selective to vicinal diols. This makes it ideal for the targeting and oxidation of carbohydrates, converting them into aldehydes and ketones. These breakdown products reside in the 5-6ppm region of the NMR spectrum, which is normally devoid of metabolites. While it is possible that NaIO_4_ may affect metabolites containing a primary amine and a hydroxyl group on an adjacent carbon atom, this oxidant does not react with the majority of other metabolites, allowing for minimal sample interference (Yuan et al., 2018).

Experiments were performed to assess whether NaIO_4_ could remove the sugars present in both IC and EC samples prepared in this study. Initially, M9 medium was lyophilised and resuspended in 50mM NaIO_4_ and then incubated for 4 hours. This was compared against a sample of M9 medium that did not undergo NaIO_4_ treatment and both samples were run on an NMR spectrometer and analysed using Chenomx software (**Figure S1**). It is evident that in the absence of NaIO_4,_ there is significant peak overlap caused by sugar contamination in the 3.2-4.5 ppm region (**Fig S1A**). Using Chenomx, we can map various peaks in the ^1^H NMR spectrum to glucose (**Fig S1B**), proving that this sugar is not removed at any stage during sample preparation. However, it can be observed that the majority of these peaks are not present in the ^1^H NMR spectrum of M9 medium when treated with NaIO_4_ (**Fig S1C**). Chenomx software was not able to match any peaks in this spectrum to glucose once NaIO_4_ was added (**Fig S1D**). Despite the significant loss of sugar peaks following treatment, a variety of relevant peaks remained in the spectrum. Chenomx allowed for the identification of 8 compounds in M9 medium following NaIO_4_ treatment (**Fig S1E**) signifying that these compounds are naturally present in the growth medium after treatment and thus can be used as a list of control compounds. Overall, this proves that NaIO_4_ is capable of removing sugar contamination in metabolomics samples, while leaving other metabolites intact.

When tested with IC samples isolated in this study, it was found that 50mM NaIO_4_ is capable of completely removing sugars from these samples and no glucose peaks were present in the spectra (**Figure S2A**). However, EC samples were shown to contain much higher glucose levels and incubations with 50mM NaIO_4_ did not significantly reduce the sugar concentration in these samples. The concentration was increased to 300mM NaIO_4_ and this showed a substantial decrease in sugar contamination, however, some carbohydrate signals still remained (**Figure S2B**). We found that we could not increase the NaIO_4_ concentration further as the oxidant became insoluble when higher than 300mM. Thus, we decided to use 300mM NaIO_4_ for 4 hours for all EC samples to maximise carbohydrate oxidation and to generate the clearest NMR spectra possible, resulting in the highest rates of metabolite detection.

### 3.4. Identification of intracellular metabolites extracted from Gram (+) and Gram (-) bacteria

^1^H NMR analyses were performed for each NaIO_4_-treated sample and all NMR spectra were processed using Chenomx software. This allowed for the detection of metabolites, as well as their quantification, relative to an internal DSS standard. An example of a ^1^H NMR spectrum displaying peaks from IC metabolites isolated from a single replicate of *E. coli* MG1655 can be seen in **Figure S2A**. In this spectrum, the peaks of the internal DSS standard are highlighted in green. This compound peaks at 4 different locations, which are 0.0, 0.6, 1.8 and 2.9 ppm. Using this standard at a defined concentration allows for the correction of the NMR spectrum and the calculation of metabolite concentration. Other notable peaks include a large peak at 4.8 ppm which corresponds to deuterium oxide (D_2_O) and a large peak at 8.4 ppm. This peak at 8.4 ppm is mapped to formate which is produced as a by-product of the NaIO_4_ interactions with carbohydrates (Yuan et al., 2018). Peaks highlighted in red signify those that are matched to compounds by the Chenomx software. However, it can be observed that not all peaks can be mapped. Also, some compounds’ peaks appear at a wide range of chemical shifts in the spectrum and it is not always possible to detect every predicted peak for a given compound.

For all IC samples (**Figure 2**), it was found that sugar contamination was removed as the Chenomx software could not map any peaks to known carbohydrates. This proves that the NaIO_4_ incubation parameters are sufficient for removing sugars present in IC samples prepared using this method. It was also observed that the 3 replicates of each strain exhibited high levels of similarity when their spectra were overlayed, highlighting the reproducibility of these methods and the suitability of this method for different groups of bacteria (**Figure 2A-D**). There were a range of compounds detected in each species (**Figure 2E-H**). For *E. coli* MG1655 (**Figure 2E, Table S1**), *K. pneumoniae* NCTC418 (**Figure 2F, Table S3**), *E. faecium* NCTC13169 (**Figure 2G, Table S5**) and *S. aureus* NCTC8325 (**Figure 2H, Table S7**), there were 15, 14, 15 and 14 compounds detected respectively across all 3 replicates. Some of these compounds like alanine, betaine, formate, isoleucine and valine are also found in the M9 medium control (**Figure S1E**). However, during the sample preparation stage for IC samples, all growth medium is removed. Thus, these compounds are also present within the cells and can therefore be deemed relevant. Despite there being some common metabolites detected across all 4 species, it can be observed that there are some unique compounds that only exist in one species. Notable unique compounds highlighted in green (**Figure 2**) are choline, glycine, imidazole and methylamine, which are all present solely in *E. coli* MG1655 (**Figure 2E**). O-phosphocholine is unique to *K. pneumoniae* NCTC418 (**Figure 2F**), while pyroglutamate and trimethylamine N-oxide (TMAO) were found to be specific to *E. faecium* NCTC13169 (**Figure 2G**). Lastly, the *S. aureus* IC extracts did not contain any unique compounds (**Figure 2H**). Excluding formate, 9 compounds were found to exist in all species (**Figure 2E-H**). These compounds are acetoin, acetone, alanine, betaine, hypoxanthine, isoleucine, leucine, phenylalanine and valine, suggesting that these compounds may be highly conserved across different bacterial species.

**Figure 2.**
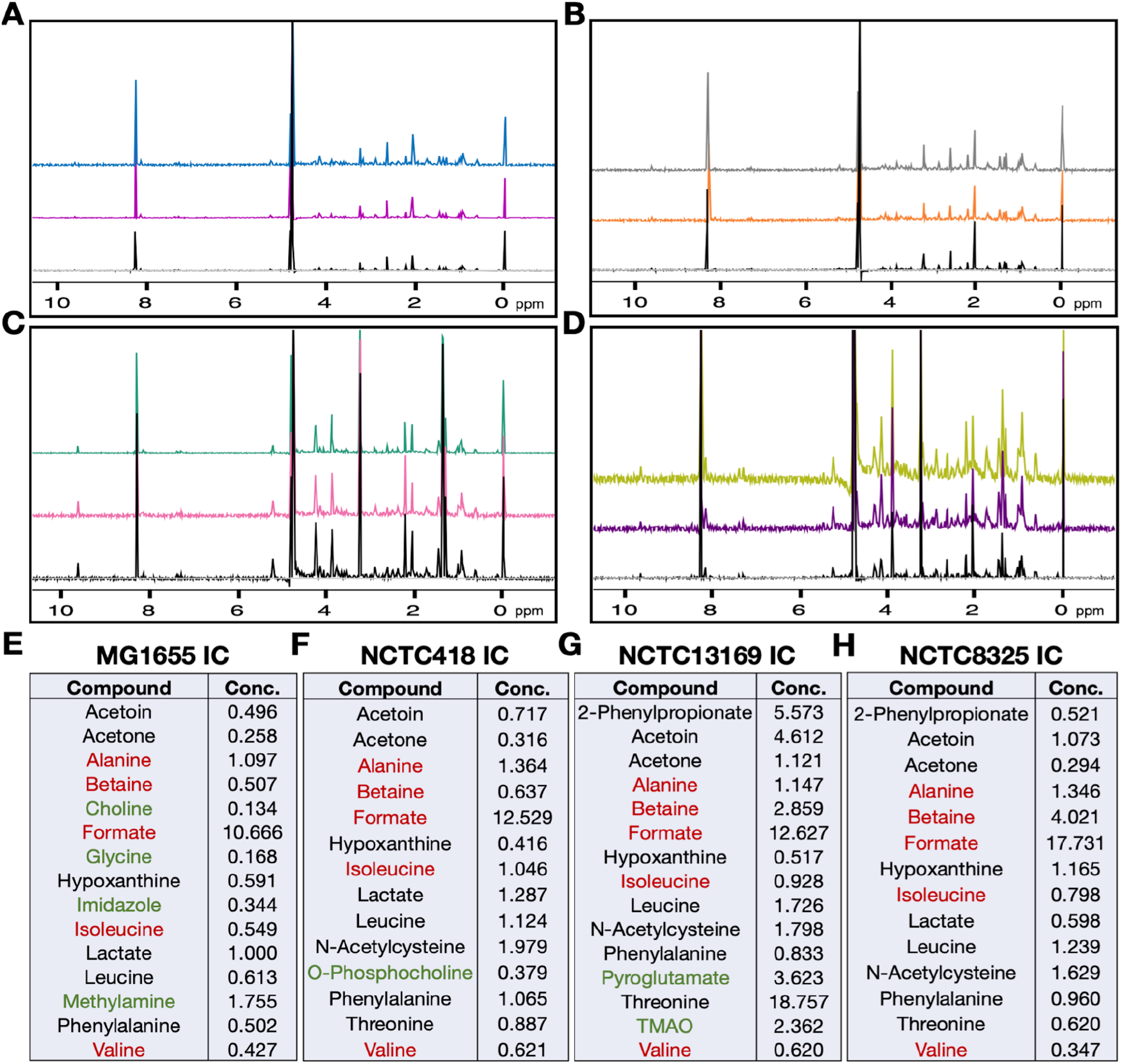
^1^H NMR spectra and identified intracellular compounds in *E. coli, K. pneumoniae, E. faecium* and *S. aureus.* ^1^H NMR spectral overlays showing 3 replicates of *E. coli* MG1655 **(A)**, *K. pneumoniae* NCTC418 **(B)**, *E. faecium* NCTC13169 **(C)** and *S. aureus* NCTC8325 **(D).** All spectral overlays were created using the Chenomx Processor module. Intracellular compounds (IC) were detected and quantified using the Chenomx Profiler module and are displayed for *E. coli* MG1655 **(E)**, *K. pneumoniae* NCTC418 **(F)**, *E. faecium* NCTC13169 **(G)** and *S. aureus* NCTC8325 **(H).** All concentration (Conc.) values are millimolar (mM) concentrations. These concentration values are the average of 3 independent biological replicates and only compounds detected in all 3 replicates are listed. Compounds in red are also found in the M9 medium control, while compounds in green are only found in one species. Compounds in black are found in more than one species.

### 3.5. Identification of extracellular metabolites extracted from Gram (+) and Gram (-) bacteria

The intensities of the peaks are considerably higher for EC compounds than for IC compounds in the example shown of *E. coli* (**Figure S2)**. The DSS internal standard peaks are detectable at 1mM concentration, which is as expected. However, they are mostly obscured due to the relative intensities of other peaks in the spectrum. This suggests that there are high concentrations of compounds secreted by cells into the surrounding growth medium. In both the IC and EC spectra, a large peak at 4.8 ppm for D_2_O is present, as is the peak for formate at 8.4 ppm, providing evidence that the NaIO_4_ reaction took place. However, it is evident that glucose contamination still remains in the region of 3.2-4.5 ppm. This suggests that the NaIO_4_ concentration is not sufficient to remove all of the glucose from the samples. Unfortunately, the NaIO_4_ concentration cannot be increased further as this causes an issue with solubility of the compound. However, the sugar concentration is reduced when 300mM NaIO_4_ is used and non-carbohydrate compounds can still be resolved and mapped in this region of the spectrum. Also, it is noteworthy to mention that there are peaks in the NMR spectra for EC samples that cannot be matched to any compounds in the Chenomx database. There are also compounds that peak in multiple regions of the spectrum and not all peaks for a specific compound may be mapped.

For all EC metabolite samples, the data was reproducible, which can be observed in the NMR spectral overlays for each species (**Figure 3A-D**). It is evident that, when compared to the 1mM DSS control, the intensities of the EC compound peaks in each spectra are considerably higher than those observed for IC compounds (**Figure 2**), suggesting that large quantities of metabolites are secreted from the cells. Unfortunately, like in **Figure S2B**, there is glucose contamination present in the spectra. However, it was possible to identify numerous non-carbohydrate compounds in these regions. Overall, the number of compounds identified in the EC samples (**Figure 3E-H**) was higher than the numbers observed in the IC samples. For *E. coli* MG1655 (**Figure 3E, Table S2**), *K. pneumoniae* NCTC418 (**Figure 3F, Table S4**), *E. faecium* NCTC13169 (**Figure 3G, Table S6**) and *S. aureus* NCTC8325 (**Figure 3H, Table S8**), there were 21, 20, 22 and 24 compounds detected respectively across all 3 replicates. Compounds found in the EC samples that are also found in the M9 medium control are alanine, betaine, formate, glycolate, isoleucine, acetate and valine (**Figure S1E**). However, aside from formate and glycolate, the remainder of these compounds were found to be considerably higher in concentration in the 20mL EC samples when compared to the concentrations of the same compounds detected in 20mL of the medium control. This suggests that these compounds are secreted by cells in quantities far in excess of the baseline levels found in the medium. There are common metabolites detected across all 4 species, as well as compounds that are unique to each species. 2-phenylpropionate, 3-hydroxyisovalerate, glycine, homoserine and thymine are all unique to *E. coli* MG1655 (**Figure 3E**). Isobutyrate and methanol were found to be specific to *K. pneumoniae* NCTC418 (**Figure 3F**). 2’-deoxyuridine, 5-aminopentanoate, carnitine and pantothenate are unique to *E. faecium* NCTC13169 (**Figure 3G**). Lastly, 3-phenylpropionate, cadaverine, gluconate, glutamate and tropate are all specific to *S. aureus* NCTC8325 (**Figure 3H**). Excluding formate and carbohydrates like fucose and glucose, 7 compounds were found to be common to all species. These are alanine, betaine, isoleucine, leucine, phenylalanine, pyridoxine and valine, suggesting that these compounds are highly conserved across different species. It can be noted that there are also certain compounds that are common to 3 out of 4 of the bacterial species, signifying that these may also be conserved across different bacterial families. These include cystathionine which is found in all species except *S. aureus*, hypoxanthine which is detected in each species, aside from *E. faecium* and both 1,3-dimethylurate and 2-hydroxyisobutyrate which are absent from *E. coli* but are detected in the data sets for each of the other species tested.

**Figure 3.**
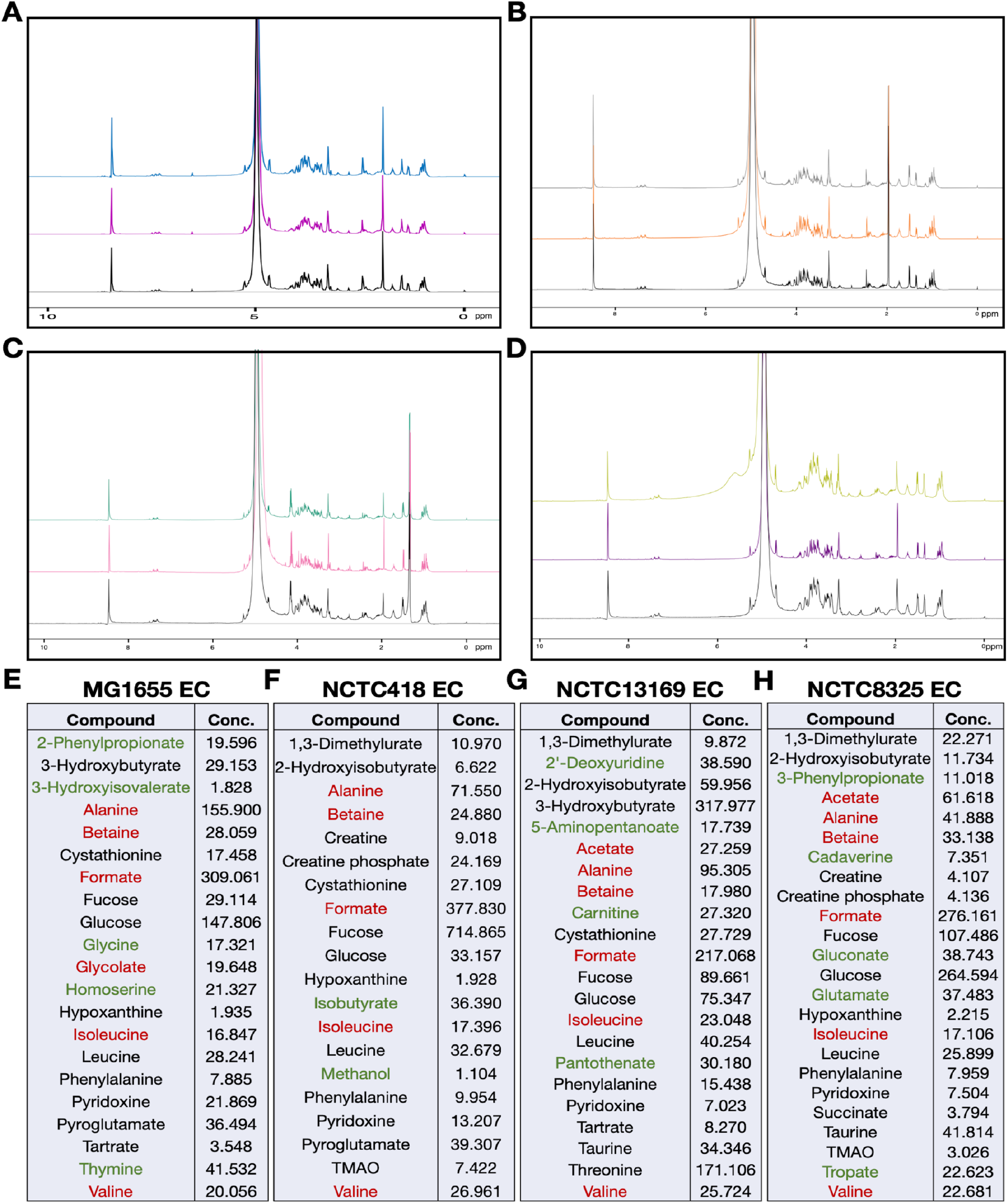
^1^H NMR spectra and identified extracellular compounds in *E. coli, K. pneumoniae, E. faecium* and *S. aureus.* ^1^H NMR spectral overlays showing 3 replicates of *E. coli* MG1655 **(A)**, *K. pneumoniae* NCTC418 **(B)**, *E. faecium* NCTC13169 **(C)** and *S. aureus* NCTC8325 **(D).** All spectral overlays were created using the Chenomx Processor module. Extracellular compounds (EC) were detected and quantified using the Chenomx Profiler module and are displayed for *E. coli* MG1655 **(E)**, *K. pneumoniae* NCTC418 **(F)**, *E. faecium* NCTC13169 **(G)** and *S. aureus* NCTC8325 **(H).** The concentrations (Conc.) listed are the values (in mM) detected in 20mL of supernatant and are the averages of 3 independent biological replicates. Only compounds detected in all 3 replicates are listed. Compounds in red are also found in the M9 medium control, while compounds in green are only found in one species. Compounds in black are found in more than one species.

## 4. Discussion

Untargeted metabolomics has been shown to be a useful tool for unravelling the complex metabolic changes that occur in bacteria in response to antibiotics and environmental stresses (Patejko et al., 2017). It has been shown that various metabolic processes can influence both drug susceptibility (Lobritz et al., 2015) and sensitivity to various stress agents (Gaucher et al., 2020; Kurake et al., 2019). Therefore, it is important to decipher the mechanisms that are utilised by bacteria in order to adapt and respond to changes in their environment. A number of drawbacks in the field of NMR metabolomics has resulted in the limited use of these techniques in microbiology research. While studies have been performed using untargeted metabolomics approaches for bacteria, they are often limited to individual species or they solely focus on isolating IC or EC compounds and not both simultaneously (Aries and Cloninger, 2020; Kumar et al., 2022; Ye et al., 2021). When both groups of compounds have been extracted in a singular protocol, it has commonly been performed using a singular species (Foschi et al., 2018; Hoerr et al., 2016). In this study, we have developed a novel untargeted metabolomics workflow that enables simultaneous isolation of IC and EC metabolites from a selection of clinically relevant Gram (+) and Gram (-) bacteria. This workflow also facilitates the detection and quantification of isolated compounds using ^1^H NMR and Chenomx software, offering distinct advantages in comparison to MS-based methods (Zhong et al., 2022). This will help simplify NMR metabolomics research in microbiology. Consequently, this will aid in the characterisation of metabolic responses to stress agents and may contribute to the identification of new targets to help combat antimicrobial resistance.

For *E. coli* MG1655, The IC (**Figure 2E**) and EC (**Figure 3E**) samples showed considerable differences in the compounds detected. The presence of unique compounds in each data set suggests that cell leakage is minimal using this protocol as more overlap would be expected if the cell membranes had been damaged prior to separation of the IC and EC fractions. This also provides evidence that certain compounds are secreted by cells into their surrounding environment while other metabolites are retained within the cell. Overall, across the IC and EC datasets, 27 distinct compounds were detected. Previously, it was shown that simultaneous IC and EC extraction from *E. coli* subjected to various antibiotics resulted in 23 IC and 20 EC metabolites detected, with it not being stated how many compounds are unique to each dataset (Hoerr et al., 2016). These numbers are slightly higher than the overall numbers detected in our study. However, this is expected as antibiotic stress likely induces the production of more unique compounds in comparison to the control conditions tested in our investigation. Another NMR study isolated IC compounds from both a wild type and a mutated *E. coli* strain, each grown to mid-log and stationary phase. A total of 31 compounds were detected across all 4 datasets (Aries and Cloninger, 2020), 9 of which are also found in the IC samples in our study, which only looked at IC compounds from one *E. coli* strain at one defined growth phase. Taken together, these data suggest that our protocol can result in the isolation of similar numbers of compounds to other workflows established for *E. coli*.

For *K. pneumoniae* NCTC418, 26 distinct compounds were detected across the IC and EC datasets. A previous ^1^H NMR study aimed to characterise the IC and EC metabolomes of carbapenemase-positive and negative *K. pneumoniae* strains in basal conditions and in response to meropenem stress. They found that 20 IC and 40 EC compounds were detected across all 59 strains used and these metabolites could be predominately grouped as alcohols, amino acids and organic acids (Foschi et al., 2018). In our study, 14 of these compounds were also found and they could be grouped into similar families. The higher number of compounds detected by (Foschi et al., 2018) is likely due to strain to strain variations across their collection of strains, as well as the antibiotic conditions utilised in the study, in comparison to our work which analysed one *K. pneumoniae* strain in basal conditions. Another recent ^1^H NMR study isolated IC compounds from various Gram (-) and Gram (+) species, including *K. pneumoniae* (Mielko et al., 2021). They tested 3 different extraction methods and found a combined total of 30 IC compounds. Of the 14 compounds detected in IC samples from *K. pneumoniae* in our study, 10 were also detected by (Mielko et al., 2021), signifying that there is significant overlap in the metabolites detected. This suggests that our method is suitable for the isolation of both IC and EC compounds from *K. pneumoniae* and can give comparable results to other validated protocols for this species.

For *E. faecium* NCTC13169, there were 29 individual compounds detected across all IC and EC datasets. Metabolomics studies on *E. faecium* are limited. NMR metabolomics has been used to identify specific bacterial species like *E. faecalis* in clinical samples based on metabolic profiling of supernatants, aiding in clinical diagnosis (Palama et al., 2016). However, as of yet, this has not been proven to be applicable for *E. faecium*. One recent study was found that utilised ultra-performance liquid chromatography tandem mass spectrometry (UPLC/MS/MS) and the advanced mass spectral database mzCloud to analyse the metabolome of *E. faecium* (Yu et al., 2020). In this study, they detected 1,320 metabolites, which is considerably higher than the capabilities of NMR metabolomics and Chenomx software. However, these data are not comparable with our workflow as the sensitivity of NMR is known to be notably lower than MS/MS methods (Kumar et al., 2022) and the Chenomx library (version 9.0) only contains information on 336 metabolites. However, despite the reduced sensitivity of NMR metabolomics, this technology is considered to be simpler, faster, more cost-effective and more reproducible than MS/MS methods (Zhong et al., 2022), making NMR highly desirable for metabolomics research. In our study, the quantities of compounds detected in both IC and EC fractions of *E. faecium* were similar to other species tested. We also found unique compounds that were only detectable in *E. faecium* samples and not in any of the other species tested. This provides evidence that our method is suitable for Gram (+) species like *E. faecium* and these results could enable future ^1^H NMR studies to be performed and compared with our baseline data.

For *S. aureus* NCTC8325, a total of 30 unique metabolites were detectable in the *S. aureus* IC and EC extracts. The IC and EC metabolomes of methicillin-resistant and susceptible *S. aureus* strains grown as biofilm and planktonic cultures were analysed by ^1^H NMR (Ammons et al., 2014). Metabolites were extracted from samples at various stages of culture, ranging from exponential growth to the late-stationary phase and samples were also taken every 24 hours, up to the 72 hour time point. Collectively, 120 compounds were identified in the supernatants, while 40 metabolites were detected in the cell pellets. Of the 30 distinct compounds detected in our study, 23 of these were also found by (Ammons et al., 2014), signifying that there is a high degree of overlap between the data sets. While the number of compounds they detected is considerably higher, this is likely due to the variety of strains used and the range of culture conditions. We would expect to see the numbers of compounds increase using our method if more conditions and strains are incorporated. Another ^1^H NMR study aimed to characterise the IC and EC metabolomes of *S. aureus* grown under aerobic and anaerobic conditions (Sun et al., 2012). After 36 hours of growth in each condition, samples were isolated and in total, 50 compounds were identified. 16 of these compounds were also found in our datasets, representing more than half of all of the identified metabolites in our study for *S. aureus*. This provides further evidence that the compounds detected using our method exhibit significant overlap with established methods for *S. aureus*. Overall, this suggests that our method is suitable for the isolation and analysis of IC and EC metabolites from the Gram (+) species *S. aureus*.

## 5. Conclusion

In conclusion, we present a novel ^1^H NMR-based untargeted metabolomics workflow, suitable for analysing the metabolomes of bacteria. This simplified method allows for the isolation of both IC and EC compounds simultaneously from a selection of clinically relevant Gram (+) and Gram (-) species. This protocol has been shown to exhibit highly reproducible results for *E. coli, K. pneumoniae, E. faecium* and *S. aureus* in control conditions. Overall, the workflow defined in this study will simplify metabolome characterisation in bacteria in comparison to current NMR-based metabolomics protocols.

## 6. Acknowledgements

This research was supported by a Science Foundation Ireland FFP award (SFI/20/FFP-A/8924_WALSH) to Prof Walsh. The NMR facilities were funded by the Science Foundation Ireland 2012 Strategic Opportunity Fund (Infrastructure award 12/RI/2346/SOF). The authors would like to thank Dr. Ciara Tierney for proofreading this manuscript.

