## Supplementary Figure S1 and Figure S2 for "Improved workflow for untargeted metabolomics and NMR analysis of intracellular and extracellular metabolites isolated from Gram positive and Gram negative bacteria"

### Supplementary information

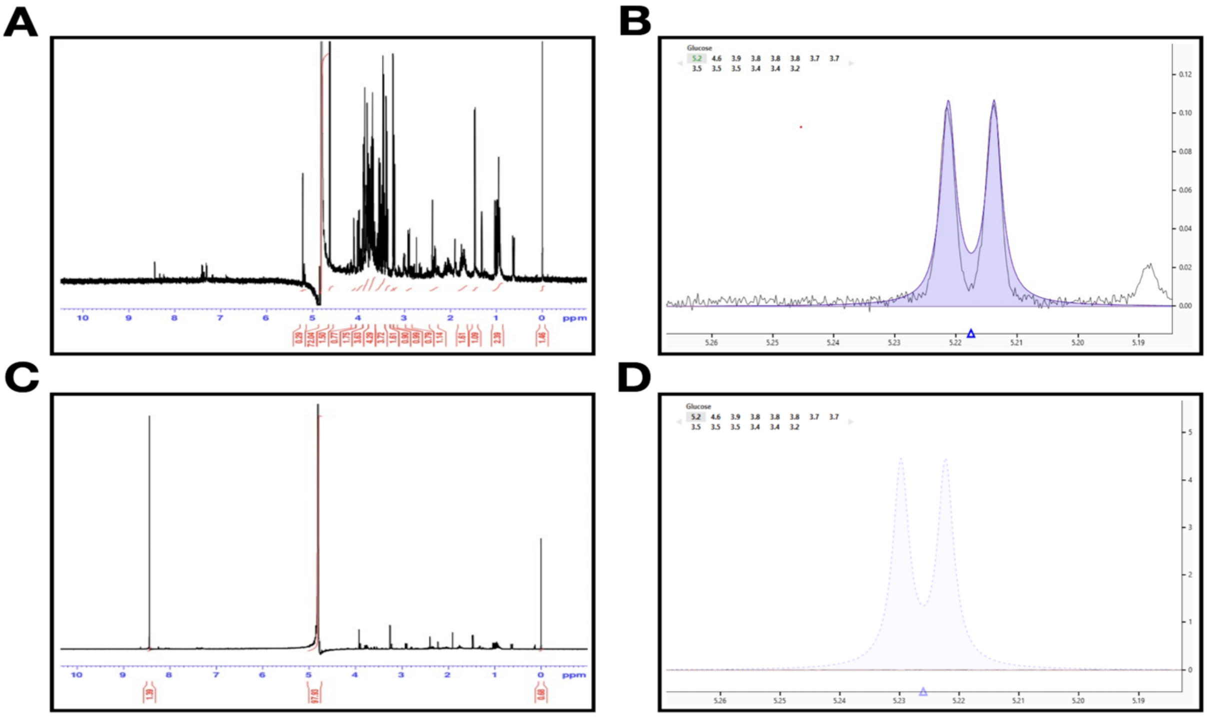


E

| Compound Name | Concentration (mM) 1ml | Concentration (mM) 20ml |
|---|---|---|
| Glycolate | 0.787 | 15.740 |
| Valine | 0.309 | 6.188 |
| Acetate | 0.449 | 8.974 |
| Levulinate | 0.207 | 4.140 |
| Formate | 19.915 | 398.294 |
| Betaine | 0.283 | 5.652 |
| Alanine | 0.705 | 14.106 |
| Isoleucine | 0.264 | 5.286 |

**Figure S1. $^1$H NMR spectra before and after $NaIO_4$ treatment. (A)** $^1$H NMR spectrum of M9 medium without $NaIO_4$ treatment. Sugar contamination can be seen in the 3.2-4.5 ppm region. **(B)** Detection of glucose peaks in M9 medium without $NaIO_4$ treatment using Chenomx software. **(C)** $^1$H NMR spectrum of M9 medium after $NaIO_4$ treatment (50mM) for 4 hours. **(D)** Absence of glucose peaks in M9 medium after $NaIO_4$ treatment. No peaks for glucose were detected by the Chenomx software. **(E)** List of compounds detected in M9 medium after $NaIO_4$ treatment using the Chenomx Profiler module. These are the compounds detected in 1mL of medium and concentration values for 20mL were scaled up accordingly.

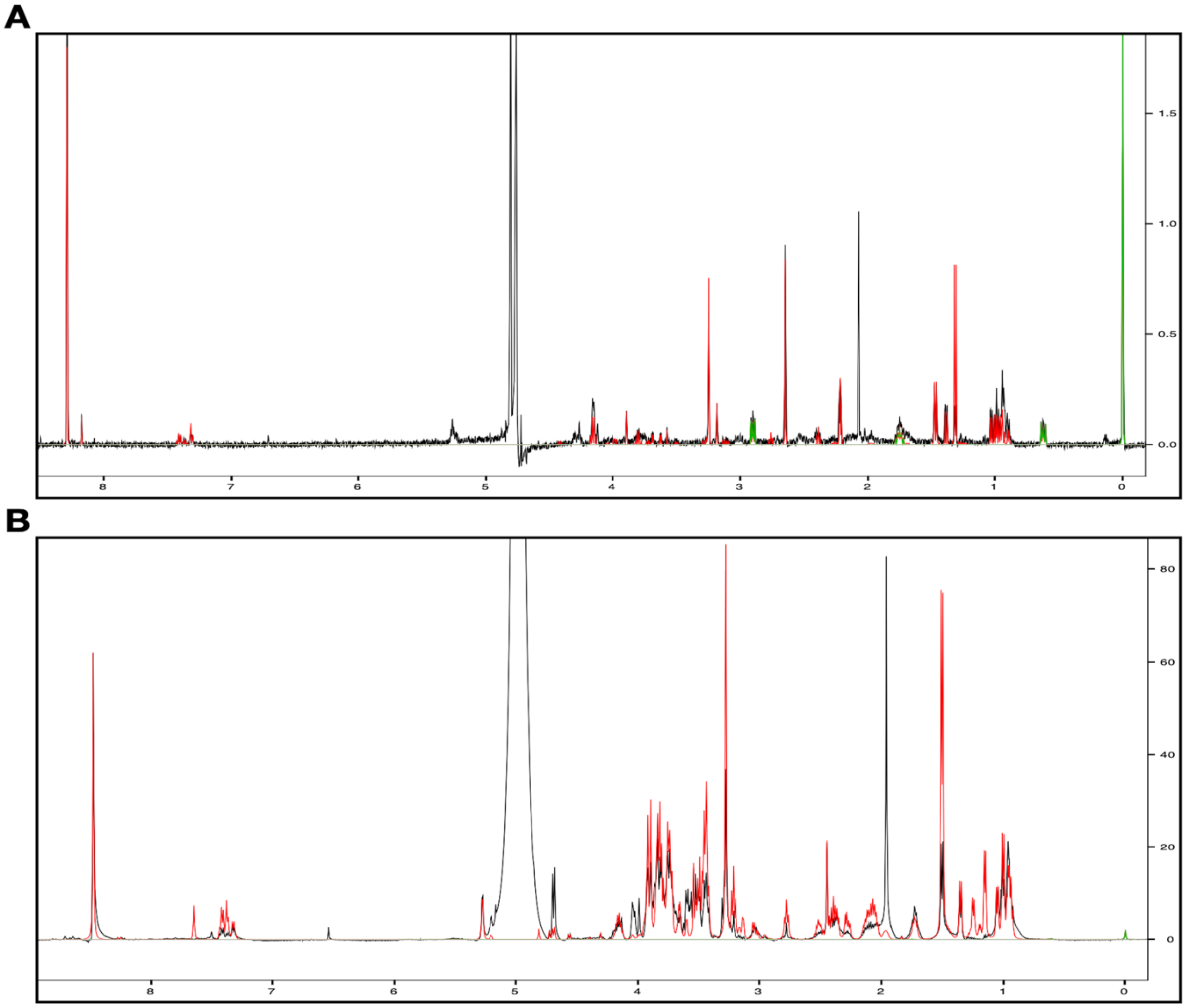


**Figure S2. Example of $^{1}$H NMR spectra for *E. coli* MG1655 intracellular and extracellular samples. (A)** $^{1}$H NMR spectrum for an intracellular metabolite extract from one replicate of *E. coli*. **(B)** Example of a $^{1}$H NMR spectrum for an extracellular metabolite extract from one replicate of *E. coli.* For both spectra, red peaks represent peaks that were matched to compounds according to the Chenomx Profiler module, while green peaks represent the DSS internal standard. Not all peaks were matched to compounds. Both intracellular and extracellular samples were treated with 50mM and 300mM $NaIO_4$ respectively, prior to loading on the NMR.
